# Cell-type specific sensing and control of firing rate statistics via channel dynamics

**DOI:** 10.64898/2026.05.01.722180

**Authors:** Andrea Ramírez Hincapié, Timothy O’Leary

## Abstract

Neurons maintain functionality through homeostatic regulation of spiking activity over extended timescales. Calcium dependent conductance expression is known to regulate mean firing rate, but this is not sufficient to ensure dynamic range in spiking activity and sensitivity to input. This raises the question of whether firing rate variance can be sensed and controlled intracellularly. Using conductance-based models, we demonstrate that time-averaged intracellular calcium dynamics inherently provide a direct readout of both the mean and variance of spiking activity. We show that calcium-based feedback regulation of membrane conductance density can therefore jointly stabilize firing rate mean and variance against input disturbances. Because tuning maximal conductances modulates the underlying relationship between rate statistics, a cell’s homeostatic response is statedependent rather than fixed. As a consequence, cell-type-specific mixtures of ionic conductances yield distinct homeostatic modalities, implying that cell type dictates homeostatic behaviour as well as spiking and integrative properties.

## Introduction

Neurons homeostatically regulate spiking activity, maintaining firing rates in a physiological range despite physiological perturbations [1–3]. Cellular mechanisms that underlie firing rate homeostasis include synaptic and intrinsic plasticity [1, 3–10]. Wide-ranging evidence has established that aver-age intracellular Calcium (Ca) concentration is a proxy for mean firing rate [11–14]. Furthermore, interfering with specific Calcium pathways is known to impair homeostatic regulation of excitable properties [15–18].

However, regulation of mean firing rate alone is insufficient for maintaining functional neural circuits. For instance, to ensure sensitivity to a fluctuating input signal, a neuron’s output variance, and possibility higher order firing rate statistics need to be regulated in addition to the mean [19– 24]. Similarly, maintenance of mean firing rate alone does not ensure appropriate patterning of activity in motor circuits [25, 26].

Regulation of higher order statistics of spiking activity requires these quantities to be sensed by biological pathways [7, 27]. Several studies have thus proposed that multiple sensing mechanisms are required to maintain spiking statistics in a functional regime [25, 28–30]. Despite several plausible models for how intracellular pathways may achieve this, there is no established cellular mechanism for doing so. This represents a major gap in our understanding of the intrinsic cellular mechanisms by which neurons adapt to continuously changing environments and input statistics.

In this work we demonstrate that Calcium concentration, averaged over appropriate timescales, implicitly senses higher order statistics of spiking activity as a result of nonlinearities in voltage gated ion channel dynamics. In other words, ion channel gating provides a direct sensing mechanism of both mean and variance of membrane potential, obviating an absolute requirement for additional intracellular pathways. This resolves an open question that has persisted since early theoretical work [31–33] and that continues to motivate experimental interrogation of large scale neuronal function in vivo, across species [34–37].

Using conductance based models derived from empirical characterisation of voltage-gated currents, we show that time-averaged intracellular Calcium concentration, ⟨[*Ca*]⟩, strongly correlates with the firing rate mean and variance under a variety of input conditions. Furthermore, the mapping between ⟨[*Ca*]⟩ and firing rate statistics is cell-type-specific, depending on the mixture of ionic currents in the membrane. Using these inferred relationships, we demonstrate a simple Calcium-based feedback loop that compensates disturbances in the input statistics to regulate mean and variance in output firing rate.

We establish conditions for which feedback control of firing rate mean and variance function reliably and show that cell-type-specific mixtures of ionic conductances lead to classes of homeo-static “personalities” that respond distinctly to changes in input mean and variance. This adds to our understanding of physiological cell type, by predicting that the dynamics of homeostatic compensation are cell-specific, and depend on both basal firing properties and channel expression profiles of neurons.

## Results

### The need for regulation of second-order firing rate statistics

To motivate the need for regulating higher order statistics of firing rates we consider the *dynamic range* problem. This is illustrated in Fig. 1a, which shows how a neuron maps input to output. To respond to changes in the input, the sensitive region of the firing rate curve must approximately match the distribution of input rates. Minimally, this requires both the midpoint, or half-maximum, as well as the width of the sloping region of the response curve to be tuned. If the distribution itself changes, the firing rate curve must adapt. Such distributional tracking has long been hypothesized to exist in sensory and other cortical circuits [38] and has since been observed in primary sensory cortices and retinal cells across species [21–24, 39].

**Figure 1.**
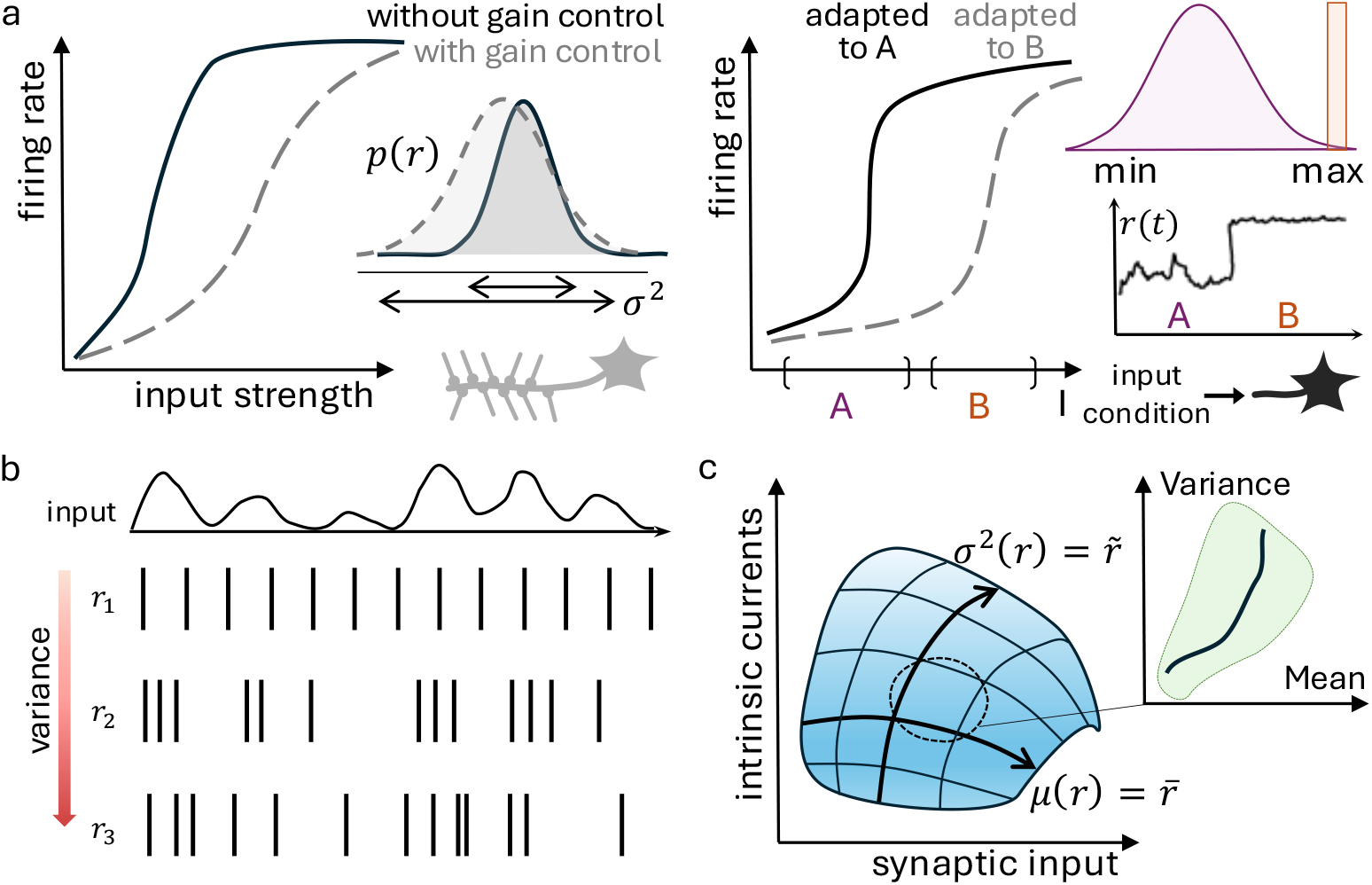
Neurons must regulate both mean and variance of their firing rate. a) Left: Cortical neurons function under high-input regimes; they would quickly saturate without gain control adjusting mean and variance of the rate distribution. Right: A sensory neuron adapted to stimulus A (purple) that does not regulate its rate distribution loses all responsiveness under new input conditions (orange). b) Mean firing rate alone is not enough to constrain the activity pattern of a neuron. Spike rasters show three degenerate solutions to a constant mean with increasingly informative variance in the inter-spike intervals. c) Tuning of various intrinsic and synaptic neuronal parameters enables the search for subspaces where target rate statistics can be satisfied. The feasible relationship between mean and variance (inset) might be constrained within a region of parameter configurations, but level sets of targets are generically transversal.

A similar problem arises for output statistics of pacemaker cells that generate neural activity in central pattern generators and other autonomous circuits [25]. A mean firing rate may correspond to various distinct firing activities. Firing rate variance, in addition to mean, can disambiguate between mean-equivalent patterns, as seen in Fig. 1b.

Both input-driven and autonomous neuronal firing properties are strongly determined by the combination of voltage-gated conductances present in the membrane. The firing rate distribution is a complex function of channel densities and synaptic input, with strong interactions between gating dynamics of different channel types mediated by membrane voltage, Calcium and other factors. As a consequence, a change in channel density of one conductance type will generically change both the mean firing rate of a neuronal membrane and the variance. This is depicted in Fig. 1c, which illustrates a 2D manifold of mean and variance level sets that depends on underlying synaptic and intrinsic conductances.

For now, there are two key points to take from this sketch: Firstly, that control of both mean and variance will generically require changes in multiple conductances simultaneously. Secondly, jointly varying firing rate mean and variance (green area) may only be feasible locally (dashed circle), where they tend to have a consistent relationship. Biologically, we posit that this local region of conductance space typifies the channel expression profile of a particular cell type. We first show that the relationship between mean and variance is inherent to neuronal intrinsic properties.

### Relationships between firing rate statistics are determined by intrinsic properties

A neuron’s firing rate, *r*, depends nonlinearly on its input history. Similarly, intracellular Calcium concentration depends not only on current events but also on past events and their nonlinear interactions. Average Calcium, taken over a time window, is therefore a (nonlinear) weighted sum, or kernel of a neuron’s recent activity history. We first show abstractly why we expect average Calcium concentration to provide a readout of multiple statistical features of firing rate, including both the mean and variance.

We approach this using the Volterra series, which is a kernel-based approximation of the input-output mapping of a nonlinear system. To see how variability in the space of (neuronal) input signals relates to variability in the space of (firing rate, or Calcium) output signals, consider *y*(*t*) = *r*(*t*) and *x*(*t*) = *I*_*app*_(*t*). The finite Volterra series for the neuronal response is

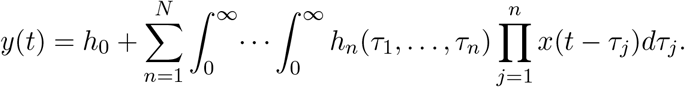

For a time window of *M* time units into the past, the integration limits become [0, *M*]. Sensible upper bounds for M are membrane time constants for mammalian central neurons, typically 20 − 100 *ms*. The *n*-th order kernels *h*_*n*_(*τ*_1_, …, *τ*_*n*_) capture nonlinear interactions of the input with itself at different time lags.

The variance and mean of the output are set by the first and second order kernels. For clarity we will assume that *h*_0_ is eliminated after centering the output and consider the two remaining leading terms, *y*_1_(*t*) and *y*_2_(*t*):

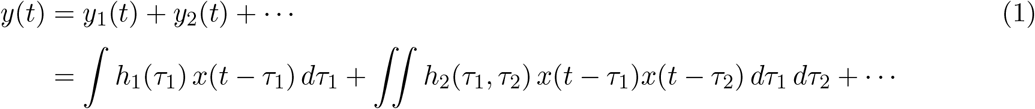

The variance of the output, Var[*y*(*t*)] = 𝔼 [*y*(*t*)^2^] −𝔼 [*y*(*t*)]^2^, can be estimated directly from these terms, noting that 𝔼[*y*(*t*)] = 0 after centering:

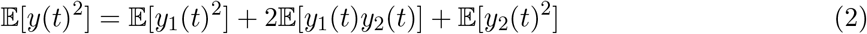

We have:

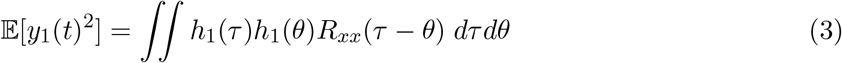

and

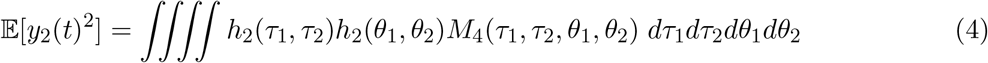

where *R*_*xx*_(*τ* − *θ*) is the input autocorrelation and *M*_4_(*τ*_1_, *τ*_2_, *θ*_1_, *θ*_2_) is the fourth-order moment of *x*.

The input autocorrelation contributes to the variance of *y* through the linear kernel in (3). Higher-order moments of the input also contribute to the variance of the output through the second-order kernels in (4), but the main contributions will be through second-order terms since the fourth-order moment of *x, M*_4_(*τ*_1_, *τ*_2_, *θ*_1_, *θ*_2_), decomposes into

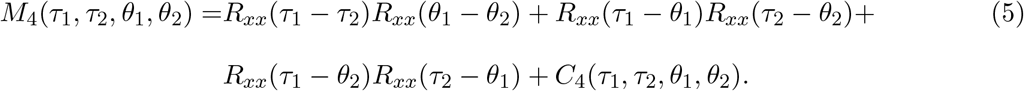

The cross terms in the expression for 𝔼[*y*_1_(*t*)*y*_2_(*t*)] in equation 2 that contain third-order moments of *x* generally do not vanish. All odd-order moments of *x* would vanish if the input was symmetric around 0 but in most realistic contexts this is not the case. Therefore, the second-order term *y*_2_ plays a dominant role in its contribution to the variance of *y*, both in the expectation 4 and in the cross-term with *y*_1_. But just like *h*_1_ shows up in cross-talk with *h*_2_ in 𝔼[*y*_1_(*t*)*y*_2_(*t*)], successive higher-order kernels will be in cross-talk with *h*_2_ when computing higher-order statistics of *y*. Therefore, as long as the the second-order kernel *h*_2_ is non-zero, it will shape higher-order statistics.

The fourth-order joint cumulant, *C*_4_, quantifies the heaviness of the tails of the distribution of *x*. For non-Gaussian distributions *C*_4_ may be nonzero. This can occur in the case of calcium concentration, which can have significant components in *C*_4_(*τ*_1_, *τ*_2_, *θ*_1_, *θ*_2_) for lags falling within the typical duration of a calcium transient.

We illustrate how all of these components behave in realistic settings using two classes of simulated conductance based neurons: a rhythmically active ‘pacemaker’ (Fig. 2a) and an input-driven spiking neuron (Fig. 2b) with (Gaussian) pink noise *x*(*t*) to model time-varying input over a finite duration. The bottom traces in Fig. 2a and b show the decay of the positive cumulant *C*_4_ as function of different lag combinations: (0, *τ*, 0, *τ*) and equidistant quadruples (0, *τ*, 2*τ*, 3*τ*). Large-amplitude events in the calcium time series at nearby times raise the expectation in

**Figure 2.**
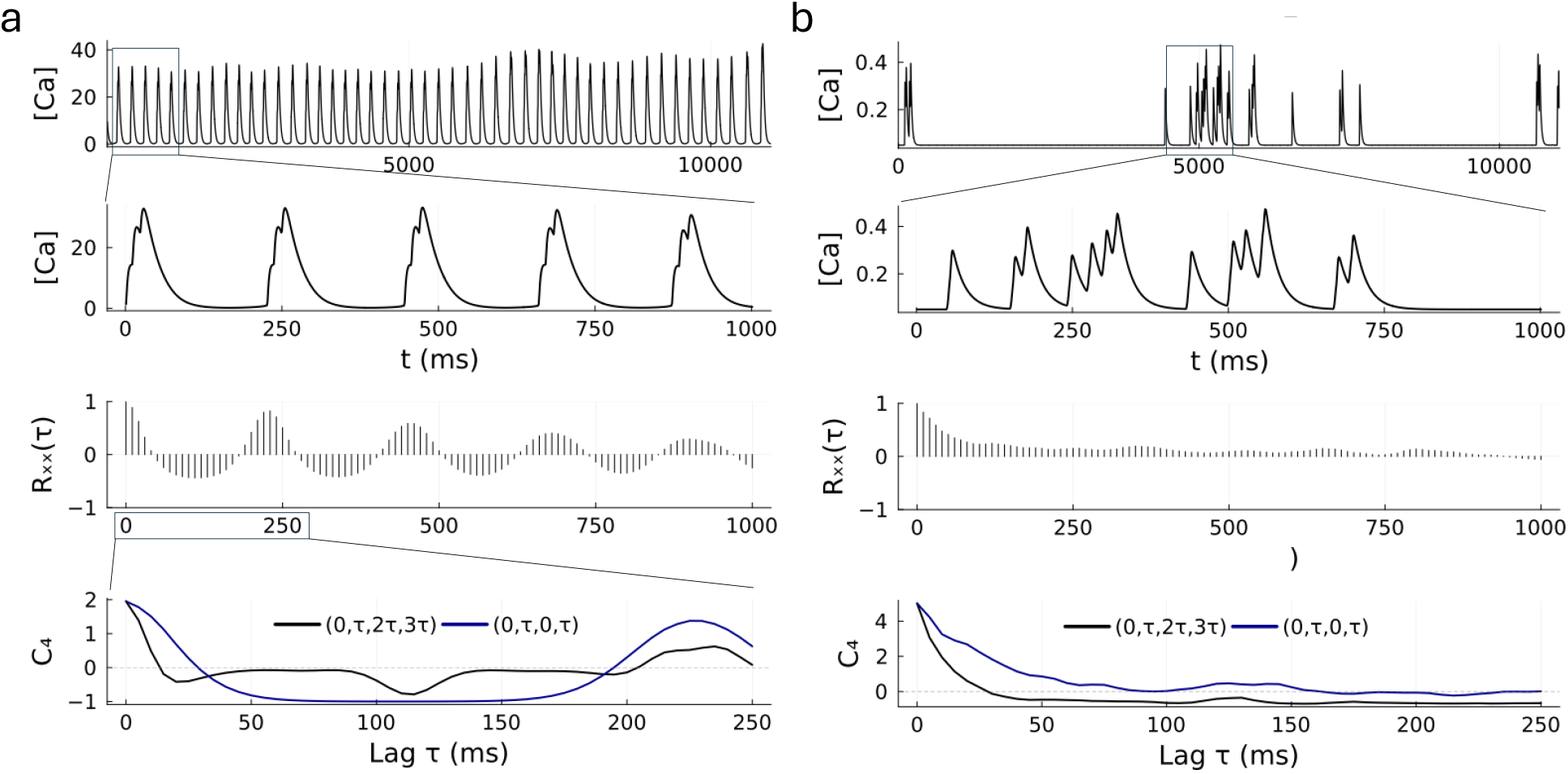
An input’s temporal structure is reported by Calcium concentration (top and inset). Fourth-order moments of the input are decomposed into autocorrelation and joint cumulant *C*_4_ (bottom two rows). Positive cumulants boost the contribution of second-order terms to the variance of a nonlinear transformation when autocorrelations are also positive. a) An active rhythmic neuron given an input with zero-mean and *σ*_*app*_ = 1.0 nA. b) An input-driven neuron given an input with *σ*_*app*_ = 8.0 nA at rheobase. In this case, it is possible to find high amplitude *τ*-distant peaks at any two time points on the lag axis plotted up to 250 ms (blue line in cumulant subplot).

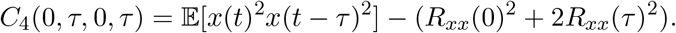

A positive cumulant in this case means that the joint squared-amplitude of *x* is larger than pairwise correlations alone would predict.

In the periodic Calcium signal (Fig. 2a) the decay of *C*_4_(0, *τ*, 0, *τ*) roughly follows the timescale of the burst envelope, approximately 40 ms. The second positive bump aligns with the bursting period, which is expected since the characteristic transient events of Calcium signals produce the largest positive cumulant if all lag arguments fall within the same burst. In the input-driven case, Fig. 2b shows that the cumulant can remain positive for a large subset of *τ* s, illustrated for both quadruple combinations, when the signal is less structured and the bursts vary in duration and frequency. In this case, it is possible to find high amplitude in the squared signal at any two time points on the lag axis plotted up to 250 ms (blue line).

The key conclusion of this analysis is that firing rate statistics are encoded in the nonlinear response kernel of a typical neuron. Furthermore, the relative contributions of mean, variance and higher order statistics are jointly constrained by the mixture of nonlinear components (voltage-dependent currents) present in the neuron. This has a significant implication for how neurons can sense and act on their firing statistics: we expect that neurons will sense different mixtures of moments via calcium concentration as a function of their membrane conductance profile and the typical input they receive. Surprisingly, in the next section we will see that these relationships are often very simple. Strongly monotonic, mostly linear, relationships exist between mean firing rate and ⟨[*Ca*]⟩ and between variance of firing rate and ⟨[*Ca*]⟩.

### Mean and variance of firing rate correlate with average Calcium

Having provided the underlying motivation for sensing mean and variance with calcium signals, we next investigated how these quantities would be sensed in large, variable populations of biophysical models. We simulated both intrinsically active and input-driven neurons across various input conditions to examine their firing rate mean and variance in relation to time-averaged Calcium, ⟨[*Ca*]⟩, which itself is computed using a cascade of two filters with plausible decay constants (see methods).

We used a stomatogastric ganglion (STG) model of a pacemaker neuron that produces a variety of periodic, endogenous spiking and bursting activity as determined by the ratios of maximal conductances in the model. We modelled an ensemble of STG neurons, initially configured in a bursting or tonic-spiking regime, with unbiased excitatory and inhibitory drive of increasing magnitude as input standard deviation, *σ*_*app*_, ranges from 1 to 14 *nA* in regular unit increments and input mean *µ*_*app*_ = 0.

Input-driven neurons, instead, were simulated with an extension of the Connor-Stevens model and were parametrised to be quiescent unless given an applied current. When input variance is ramped the input mean matches their rheobase.

Figure 3a plots mean and variance of individual STG neurons as functions of ⟨[*Ca*]⟩. Monotonic, mostly linear relationships are found. The signs of the two correlations, between ⟨[*Ca*]⟩ and mean firing rate

**Figure 3.**
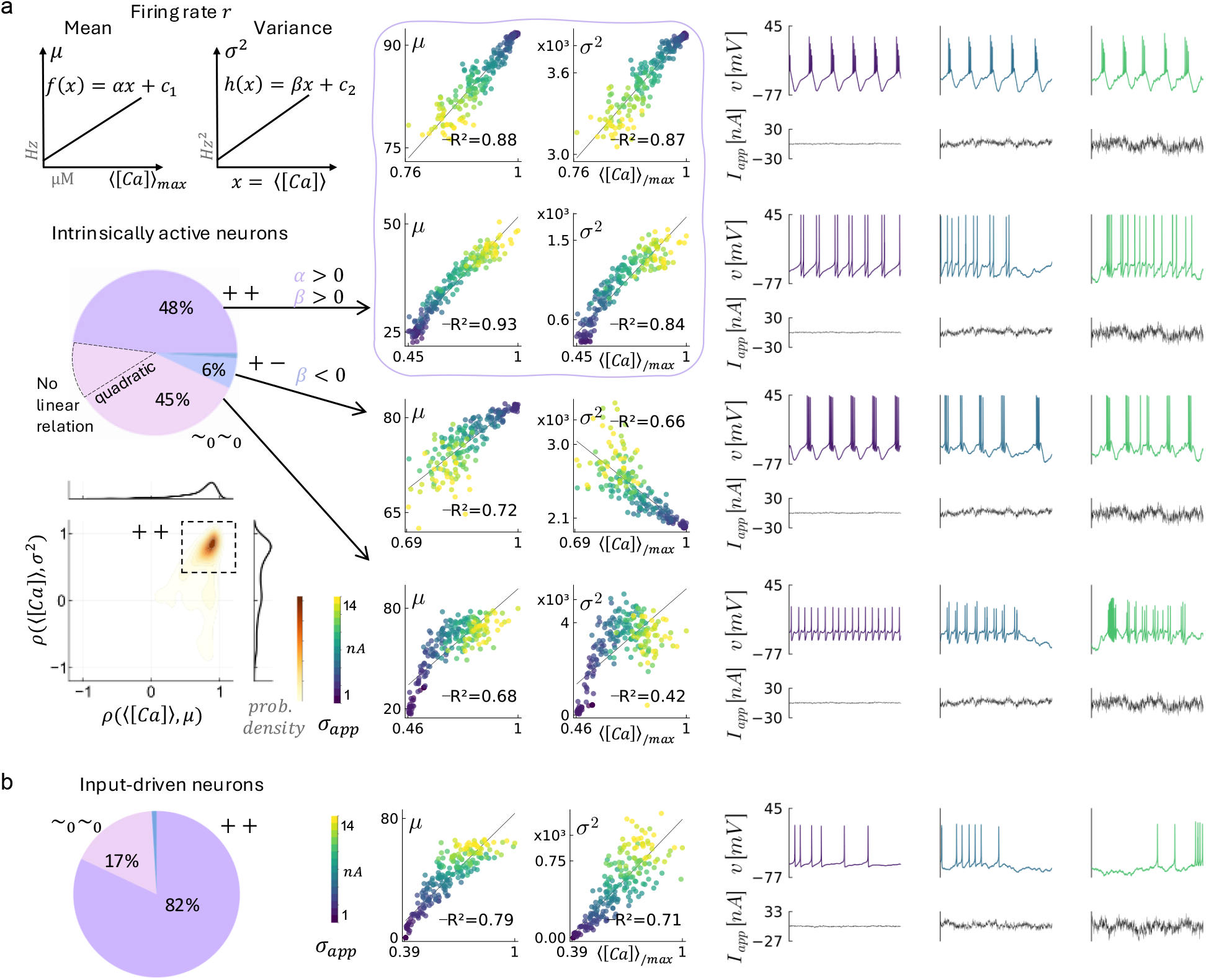
a) Top-left: Pairwise correlates of mean [Hz] and variance [Hz^2^] of firing rate with average Calcium concentration. Right: Examples of individual intrinsically-active neurons with distinct sensing modalities. Voltage traces (1 s) colour-coded as input variance increases, see *σ*_*app*_ scale bar. Bottom-left: Percentages of significant (thresholded) classes in the ensemble and marginal histograms and 2D density of correlations. Note: one fourth of the neurons in ++ class show a decrease in rate statistics as *σ*_*app*_ increases (downward trend). The x-axis is normalized to maximum ⟨[*Ca*]⟩; values for neurons are 17, 11, 19, 8 *µM* from top to bottom. b) Input-driven neurons have predominantly responsive types. Maximal Calcium is 0.22 *µM* for the example neuron on the right.

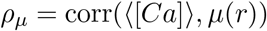

and between ⟨[*Ca*]⟩ and variance of firing rate

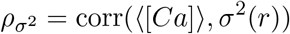

define pairwise correlation classes 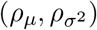 as follows. A correlation *ρ* is considered significant if |*ρ*| *>* 0.5, otherwise it is classed as a weak correlation and denoted ~_0_. Both correlations are significantly positive (++) in 48% of the neurons in the ensemble (*N* = 1080, pie chart in Fig. 3a). The mean correlates negatively with ⟨[*Ca*]⟩ only in 7% of all cases, and only in 1% of these the correlation is significant. There is no clear distinction between classes with regards to the initial firing pattern.

The top-right panel of Fig. 3a illustrates two examples of neurons with positive correlations, but with different responses to increasing *σ*_*app*_ (color coded). A neuron in class ++ may respond to increased input variance with an increase in Calcium, or the opposite effect entirely, *i*.*e*. Calcium decreases. After inspection of the members of the class with the upward trend, we find that these neurons add spikes to nominal bursts, thus increasing mean rate and mean Calcium. An increase in the number of spikes may lead to a larger repertoire of patterns if spikes cluster in or out of packets, and therefore lead to high variance too.

Increasing the input variance has very little effect on ++ neurons with the downward trend; these are strong pacemakers. As input variability increases, so does the chance of encountering inhibitory inputs that prevent spikes. Thus, while input can perturb bursting dynamics (e.g. offset them in time) and cause a decrease in mean firing rate, the intrinsic properties of a burst are relatively input-independent once a burst is initiated. Likewise, the bursty neurons in class +− are all strong bursters. As input variance drops, these neurons fall back on their regular bursting pattern which has the lowest variance at the lowest *σ*_*app*_. Similarly to the previous class, mean decreases with increased input variance but the bursting dynamics remain somewhat robust.

Around 45% of neurons in the STG population show less well-defined correlations. However, the lack of a linear relationship, or a weak one, does not exclude the possibility of nonlinear relationships. The dashed slice of the pie chart in Fig. 3a indicates that in around 20% of this subset, the relationship between mean rate and ⟨[*Ca*]⟩ is better described by a quadratic model, where the criterion for a good quadratic fit is *R*^2^ *>* 0.7. Only a handful of cases exhibit good quadratic fits between rate variance and ⟨[*Ca*]⟩ (not shown). However when the data points fan out we cannot rule out heteroscedasticity, a phenomenon where the variance of the residuals increases as a function of the deterministic variable, ⟨[*Ca*]⟩.

The vast majority of neurons in the input-driven ensemble (*N* = 588, Fig. 3b) show pairs of positive correlations with an upward trend. Naively, one expects that mean and variance both increase as input variance is ramped, since these neurons initially receive the minimal current required to elicit a spike, and so larger deflections from a cell-specific baseline drives neurons over their spiking threshold more often.

In summary, many neurons in a variable population have relatively simple relationships between average calcium and the mean and variance of their firing rate. To simplify our results in what follows, whenever approximately linear relationships hold among these quantities, we will express them as affine functions of *x* = ⟨[*Ca*]⟩

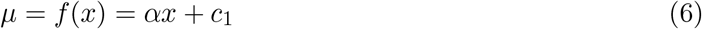

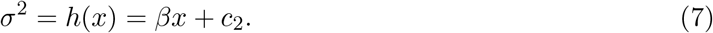

The shorthand *µ* = *µ*(*r*), *σ*^2^ = *σ*^2^(*r*) is introduced to denote mean and variance of firing rate. The cell-specific coefficients *α, β* are positive in approximately 50% of active neurons and in 82% of input-driven neurons. Next, we show that these functions are not static.

### New input statistics reveal different correlations

In networks that need to self-organize from an initially inactive or hyperactive state, a homeostatic response could achieve target activity patterns [40]. But what are the achievable rate statistics? That is, how much are mean and variance constrained as a pair? We asked if the pair *f, h* constitutes a fixed mode of a (joint) sensing mechanism of rate statistics in a cell. To assess the fidelity and variety of of sensing modalities, we examined whether model STG neurons responded differently to changes in input statistics.

We simulated input currents with increasing mean *µ*_*app*_ ranging from 1 to 10 *nA* and fixed *σ*_*app*_ = 2. The pie chart in Fig. 4 reveals a new picture for the pairs *ρ*_*µ*_, 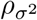under this protocol. The ++ and +− classes found in the previously are present again, but we also find a number of negative correlations in both *ρ*_*µ*_ and 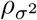. The neurons that fall in the −− class have a mean rate that is consistently bounded between 30 ± 5 Hz (top row). The firing pattern generally transitions from three-spike bursts to doublets, suggesting a stereotypical adaptation.

**Figure 4.**
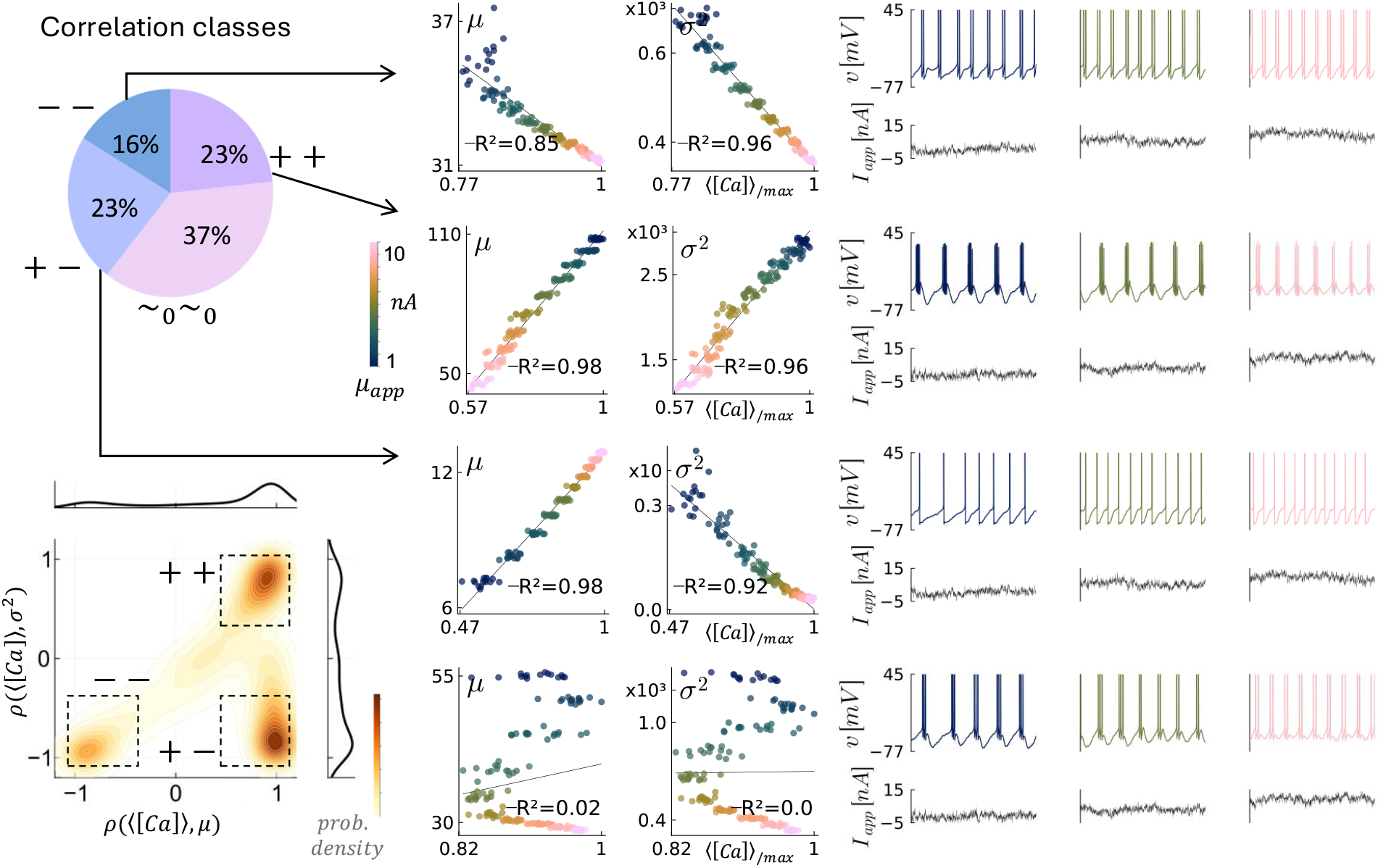
Pairwise correlates of mean [Hz] and variance [Hz^2^] with ⟨[*Ca*]⟩ as input mean *µ*_*app*_ increases. Left: Joint density of correlations across the ensemble. A large percentage of neurons in class −− becomes apparent when *µ*_*app*_ increases and *σ*_*app*_ is fixed. Right: Functional relationships and voltage traces (1 s) for individual neurons in each class, colour-coded by input mean. Non-linear relationships may be more complex than a quadratic form, as suggested by S shaped plot in the bottom row. Maximal Calcium normalized by 18, 14, 6, 12 *µM* from top to bottom.

Neurons with the highest correlations class +− are representative of fast tonic spikers which increase their firing rate as input mean increases while changes in variance are negligible (Fig. 4 third row). The same is true for the ++ sub-class with upward trend (not shown). In contrast, neurons in class ++ with a downward trend span a large range in both firing rate mean and variance.

What do these relationships mean for a pacemaker neuron? In motor systems, it is particularly important to reduce motor-neuron and muscle noise variability since a smooth contraction, as opposed to uncoordinated muscle twitches, requires pools of active neurons with high firing rates [41]. The high energetic cost of an action potential emphasizes the need for neurons that innervate shared muscle fibers to collectively reduce variance in their firing rates. If variance of firing rate is always in a fixed relationship to the mean then Calcium is an indicator of the desired firing statistics and hence of the bursting pattern. Indeed, bursting can be identified just by the combination of a high mean and non-zero variance.

The idea of computational homeostasis is relevant here as it suggests that, if the goal for a neuron is to preserve *function* then, ultimately, it is not only firing rate driving homeostatic regulation but also encoding properties, excitability and synchronization properties among others [29]. In the STG network, for example, it is relatively clear that tri-phasic bursting patterns that support digestive function are robust to various neuromodulatory conditions. From this viewpoint, cell-types encompass not only nominal behaviour and ‘steady-state’ properties (in sum, the characteristic computation a neuron performs under normal circumstances) but also the homeostatic response to perturbations. This suggests that suitable targets for the regulated variables must arise from the interaction of various physiological control loops.

The two different protocols on input statistics shed light on the fact that class identity is not necessarily shared across conditions. Figure 5 shows that a neuron changes its correlates of rate statistics, as functions of ⟨[*Ca*]⟩, according to the type of input disturbance. This shows the important role of synaptic input in uncovering different kinds of behaviour. Even though there is no explicit adaptation of neuronal parameters in this setting, this apparent adaptive behaviour is due to the interaction of the neuronal nonlinearities with the input statistics. When adaptation of neural computation must happen on a fast timescale (for example cortical neurons adjusting the slope of their fI curve to the variance of background input) the source of this adaptation is mainly in the intrinsic nonlinearities of the system and how these interact with the input [42].

**Figure 5.**
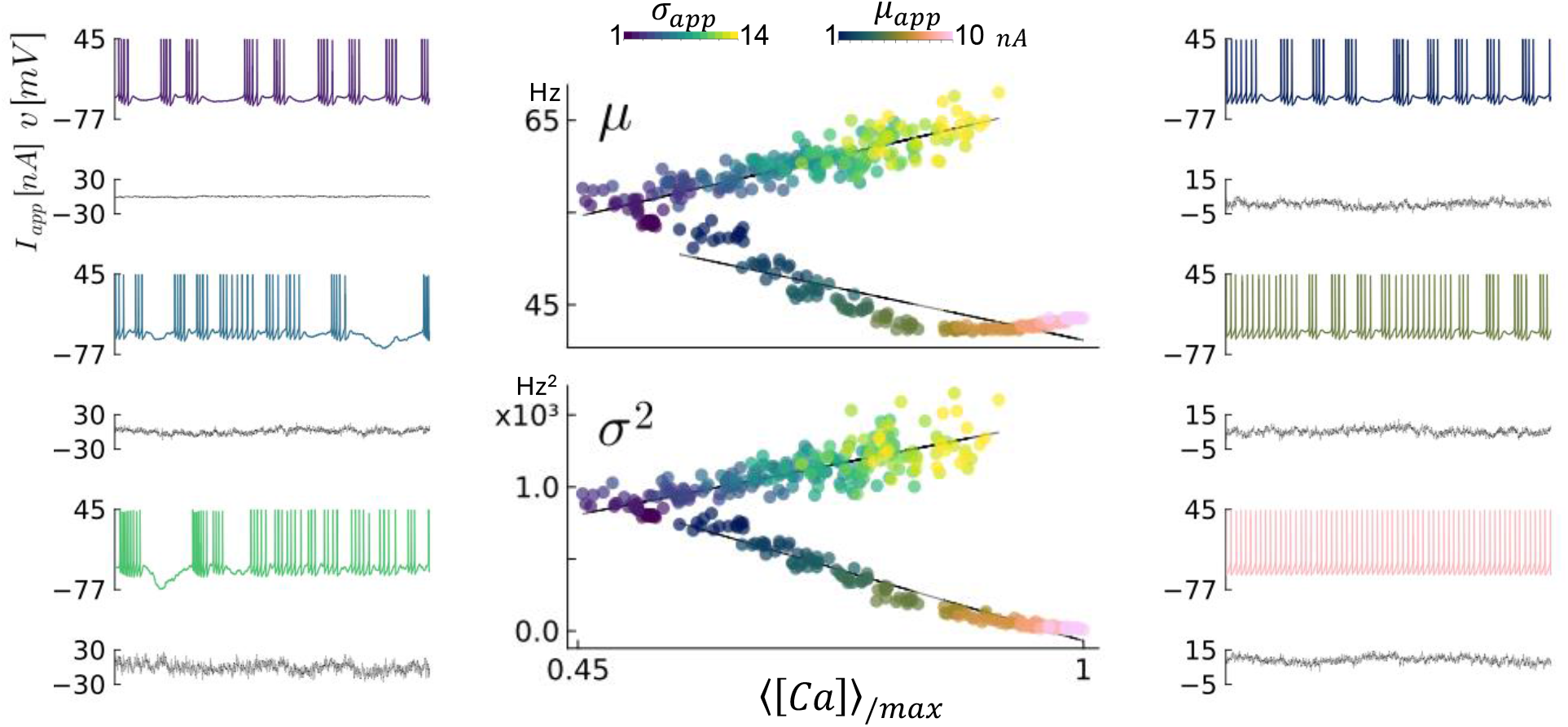
Two different ramping protocols on input conditions elicit differential responses in a neuron with the same underlying conductances. Center: Two branches of sensing correlates, *ρ*_*µ*_, 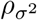, are uncovered depending on the input encountered. Voltage traces (1 s) on either side show the response to increased input variance (left) or mean (right). Maximum average calcium for x-axis normalization is 32 *µM*.

### Correlations are modulated by changes in conductances

Neurons are subject to evolving input statistics just as much as they are to evolving neuromodulatory signals. Channel densities can change considerably under different modulated states, with up to 6-fold variations in maximal conductances leading to different activity patterns [43]. Figure 6a shows that changes in a handful of conductances modulate the correlations to different signed combinations. A neuron in class ++, initially in a tonic state, is modulated to class +− with an increase of the slow Calcium conductance and a decrease in T-type Calcium conductance, 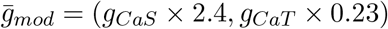.

**Figure 6.**
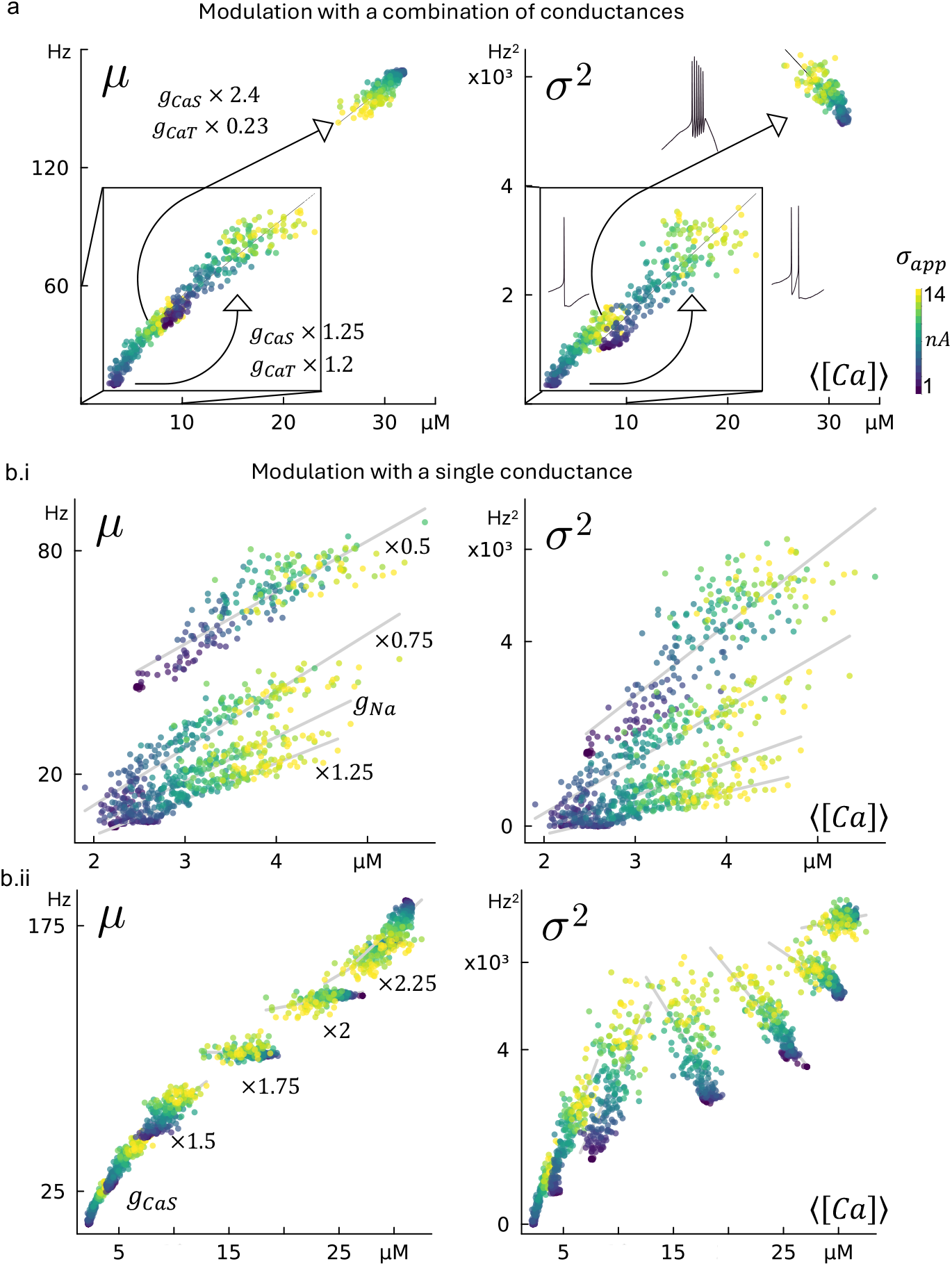
Correlation relationships depend on the modulated state of a neuron. a) Modulation in a subset of conductances changes the firing pattern and therefore the relationships between rate statistics and ⟨[*Ca*]⟩. b) A single conductance is scaled incrementally by a factor of 0.25. Sodium conductance scaled from 0.5 of its nominal value and S-type Ca conductance scaled from its nominal value.

The same neuron is modulated to different 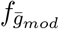 and 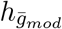 within the ++ class (box inset, closer to the initial state) through another rescaling of these two conductances. We identify *g*_*CaS*_ as one of the conductances most relevant for modulation between classes since bifurcations to bursting patterns often arise from the combination of Calcium and Ca-activated potassium currents that provide slow excitability and can drastically alter rate distributions.

Figure 6b shows that scaling along individual rays of conductances induces smooth geometric transformations (translation and rotation) of *f* and *h*. When sodium is modulated, the relationships are shifted downwards but otherwise remain qualitatively similar. This is expected since sodium is chiefly contributing to the generation of the action potential but minimally to the spiking properties. However, when S-type Calcium is increased, the nominal tonic spiking behaviour switches to bursting and this changes the direction in which the linear relationships are oriented with respect to the input.

Having a model for how neurons might sense their rate mean and variance in the pair (*f* (*x*), *h*(*x*)), we have shown that the link between the two is dependent on the modulated state of the neuron’s conductances 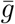. We can therefore blur the distinction between states of a neuron and neuron ‘identity’ in the ensemble. Conceptually, this state-dependence means there is a continuum of 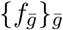 and 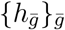. How these two change *as a pair* determines the feasible rate statistics. In principle this kind of package-deal reduces the feasible set to a one-dimensional family parametrised by 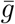, but it greatly simplifies the control of rate statistics. Next, we show how these intrinsic constraints can be leveraged for the control of firing rate statistics.

## Implications for control of rate statistics

If a neuron’s identity determines *f* and *h* then, intuitively, the reachable combinations of target rate statistics (*µ*^*∗*^, *σ*^2*∗*^) depend on the specific neuronal parameters (synaptic and intrinsic conductances in Fig. 1c). But a mixture of plasticity and homeostatic mechanisms are generally at play in any neuron, so its ability to move in this parameter space suggests there may exist a state vector 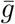 such that (*µ*^*∗*^, *σ*^2*∗*^) are in the image of 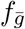 and 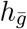. In this case it should be possible to control towards *µ*^*∗*^ and *σ*^2*∗*^ by finding *x*^*∗*^ such that 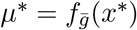 and 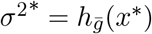.

A controller that uses ⟨[*Ca*]⟩ as the controlled variable and has two target outputs, *µ*^*∗*^ and *σ*^2*∗*^, can only satisfy the targets if they are in a certain relation to one another. As long as the function *h* is invertible, the variance can be written in terms of the mean

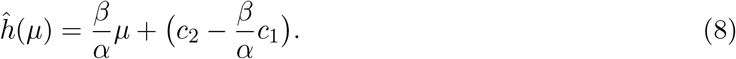

If the target in variance is *σ*^2*∗*^ = ĥ(*µ*^*∗*^) then a simple integral controller in negative feedback with the correlates of rate statistics can achieve perfect regulation [44].

Consider the controller equations

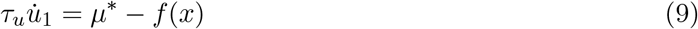

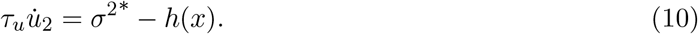

The control variables *u*_*i*_ are neuronal parameters that close the feedback loop around Calcium (see eq. 17). It is impossible to explicitly account for the myriad cellular mechanisms that act on Calcium [45, 46]. However, a biochemical realization of an integrator, one of the simplest regulation schemes, is straightforward when the rate of loss of the signal is independent from the rate of accumulation, which is the case for channel proteins in the membrane with a constant degradation rate [28]. Therefore, ion channel conductances are the natural actuators *ū*. Fig. 7a explicitly shows the dependence of sensors *f, h* on neuronal parameters *ū* in the plant, which takes in the adapted *ū* from the controller output.

**Figure 7.**
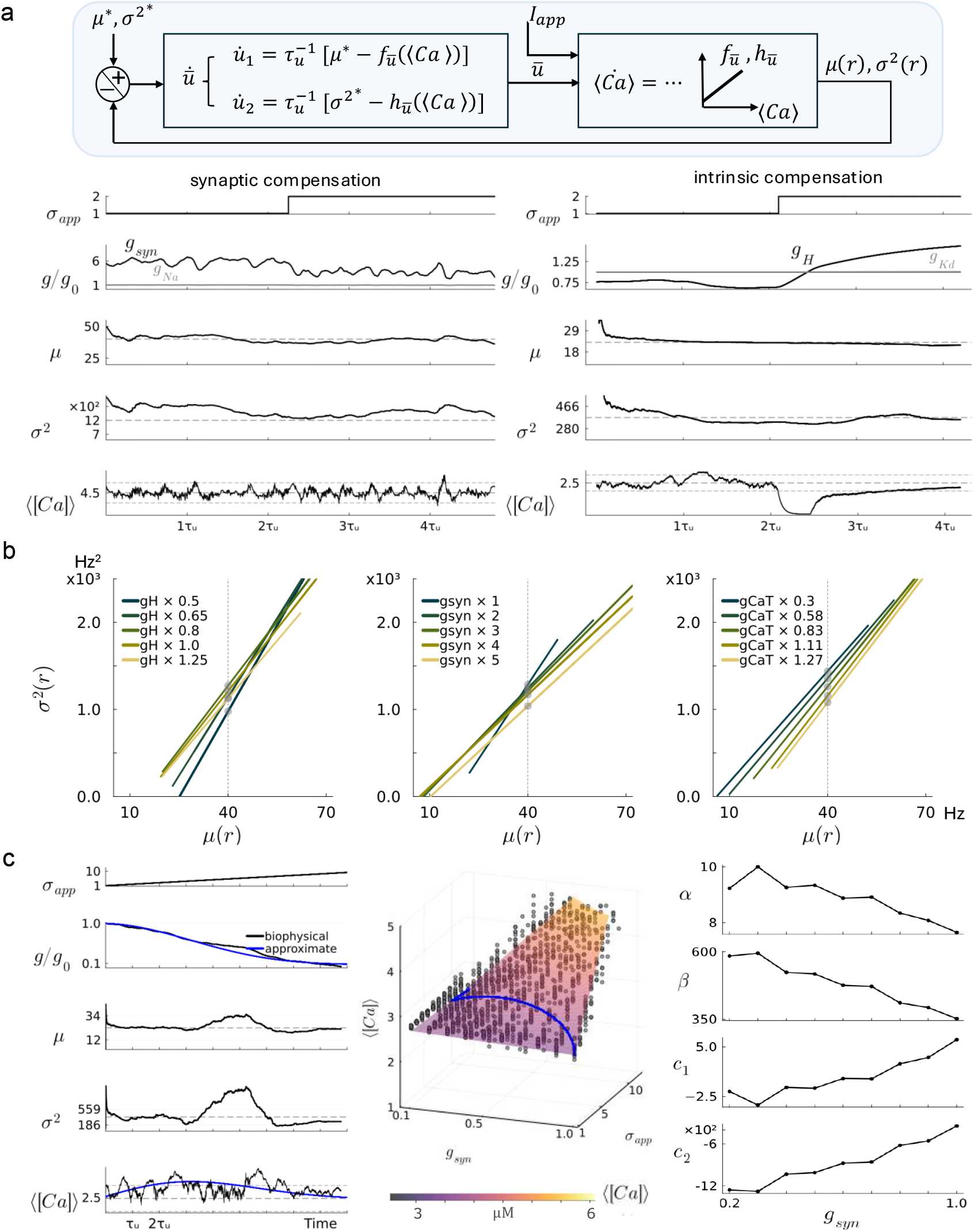
Integral control jointly regulates rate statistics. a) Control using negative feedback on estimated rate statistics (top diagram) is implemented with synaptic and intrinsic conductances. Conductance evolution and running statistics in the presence of a step disturbance in *σ*_*app*_. Guide-lines at ±25% of target ⟨[*Ca*]⟩. b) The implicit relationship ĥ is deformed due to the control action but may be more tightly preserved for certain conductances. c) Comparison of control with full biophysical model and simpler model *x*(*g*_*syn*_, *σ*_*app*_), with *γ*_0_ = 2.7, *γ*_1_ = −0.59, *γ*_2_ = −0.05 and *γ*_3_ = 0.45. Coefficients of the sensing modes vary smoothly with the control parameter *g*_*syn*_.

When *τ*_*u*_ is large enough (regulation of conductance expression is on the order of minutes), the control variables slowly integrate the two error signals. Applying the transformation *ĥ*

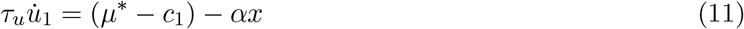

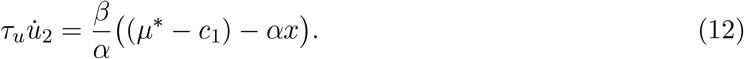

So, starting from small initial conditions, there is really only one requirement for the controllers to achieve perfect control, which is

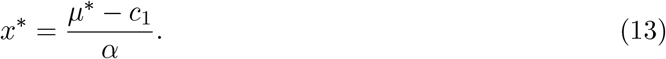

Equation 13 requires that, if *α >* 0 then *µ*^*∗*^ *> c*_1_ must hold in order for *x* (concentration) to remain positive. Conversely, if *α <* 0 then *µ*^*∗*^ *< c*_1_. This implies that sensing modes with positive (resp. negative) linear coefficients *α, β* have a lower (resp. upper) bound in their firing rate mean and variance.

System 11-12 is implemented with intrinsic and synaptic conductances. Experimental observations on synaptic scaling support the choice of the latter [2]. Figure 7b shows the controller compensates a step increase in input variance when *u*_1_ = *g*_*Na*_ and *u*_2_ = *g*_*syn*_. A target *µ*^*∗*^ = 40 Hz gives an implicit target in variance *σ*^2*∗*^ = 1202 Hz^2^ and, using equation 13 for this particular active neuron with known coefficients *α, β >* 0, a target in average Calcium *x*^*∗*^ = 4.5 *µM*.

A rolling window of size *w*, on the order of *τ*_*u*_, is used to compute online rate statistics *µ*(*r*(*t*)), *σ*^2^(*r*(*t*)) for the longer time course over which controller dynamics evolve. It is therefore unsurprising that average Calcium fluctuates around target since the controlled variables *µ*(*r*(*t*)), *σ*^2^(*r*(*t*)) are dynamically sensed as the input disturbs the system. It can be seen that one of the control variables, *g*_*Na*_, remains fixed, corroborating the system has one extra degree of freedom. Meanwhile, the synaptic conductance opposes the change in input variance. The synaptic conductance *g*_*syn*_ can also maintain average Calcium when the input is a square pulse (supplemental Fig. **??**).

A different choice of control variables, *u*_1_ = *g*_*H*_ and *u*_2_ = *g*_*Kd*_, also achieves successful compensation of rate statistics. In Fig. 7c the target is *µ*^*∗*^ = 23 Hz. In this case *g*_*H*_ alone is responsible for the control action. Panels a and b show that both intrinsic and synaptic conductances can homeostatically maintain mean firing rate, but the variance in firing rate is less tightly maintained around target (dashed line) in panel a. The difference lies in how the nominal correlates, and consequently the implicit relationship between mean and variance, *ĥ*, is deformed as the effective control variable is tuned by the controller. Fundamentally, the pair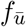, 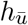 produces inaccurate estimates of rate statistics as soon as *ū* changes.

Figure 7d shows how various conductances have increasingly disruptive effects (from left to right) on sensor correlates. Contrasting *g*_*Kd*_ and *g*_*CaT*_ shows the former is a better candidate among the conductances of the STG model to retain the blueprint of nominal correlates. Maintaining a target value in average Calcium concentration through regulation of *g*_*CaT*_ alone is entirely possible, but *ĥ* is not tightly preserved. Thus, if Calcium conductances are used as control variables, a bias is introduced in the error terms and the true rate statistics cannot be reliably matched to the targets despite constant ⟨[*Ca*]⟩ (supplemental Fig. **??**). So for the controlled system to remain self-consistent, the control action *ū* (*t*) should interfere minimally with the nominal correlates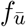,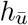.

A simplified system is implemented to validate that the control of rate statistics really is based on controlling average Calcium towards *x*^*∗*^. If the sensing modes *f, h* are sufficiently smooth as ū changes, then the stability of the rate statistics can be evaluated through the stability of ⟨[*Ca*]⟩. In the full biophysical model, average Calcium is costly to compute since it requires simulation of the internal dynamics of our conductance-based models. But the idea depicted in the control block diagram is that we just need a good approximation of the dependence of average Calcium on the effective control variable *u* and on the disturbance.

We fit a linear model to *x* = *x*(*g, σ*_*app*_) with data acquired over a trajectory of *g*_*syn*_ by repeating the input variance protocol at sampled *g*_*syn*_ values. The model is

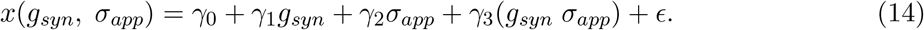

Using the predicted value of *x* from 14, the simplified control of rate statistics is

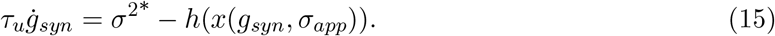

This simplified system qualitatively reproduces the results of the full biophysical model (compare black and blue trajectories). We conclude that all that is required to control rate statistics is the cell-specific information on how average Calcium responds to intrinsic and synaptic parameters, which is essentially how the sensing modes depend on input and conductances.

## Discussion

Experimental measurements across nervous systems reveal tight regulation of firing rate statistics. Motivated by experimental observations, previous computational works have proposed models of plasticity mechanisms, both intrinsic and synaptic, to regulate neuronal firing based on mean and variance of firing rate [33, 47, 48]. An early, influential model showed how intrinsic currents could be tuned to minimise the Kulback-Liebler divergence between a target firing rate distribution and the output of a detailed biophysical model [19]. Nevertheless, these models have largely addressed the problem from the perspective of actuation — specifying how plasticity rules drive firing statistics toward a target — rather than from the perspective of estimation, namely how a neuron might internally represent or infer its own firing statistics.

The excitable and integrative properties of a neuron depend on the combination of ion channels expressed in its membrane. The notion of cell type is strongly linked to channel expression [40, 49, 50] providing the biological basis for cell-type specific physiological properties, and a means of reliably encoding functional properties in a wider circuit. We have shown a novel role for channel dynamics as an internal sensor of the input-output properties of a neuron. Most notably, in the same way that combinations of channel densities determine excitable properties, different combinations of channels suppress or amplify changes in firing statistics, providing a tunable sensor of complex spiking activity.

These findings suggest two major shifts in perspective on homeostatic plasticity and its relation to cell type. First, we propose expanding the notion of neuronal cell type beyond conventional properties such as morphology, neurotransmitter composition and basal firing properties to encompass the class of signal statistics that a neuron can sense and adapt to. If this broad hypothesis is valid, experiments should reveal that certain types of neurons will adapt to specific changes in firing rate statistics that fail to cause significant homeostatic adaptation in other cell types. For example, we would predict that rhythmically active motor neurons have sensing tuned to rhythmic input and output. Early work provided some evidence of this in dissociated neurons from the Lobster Stomatogastric ganglion [51]; it would be intriguing to revisit this basic finding in the context of non rhythmically-active, or phasically active neurons and circuits.

Second, our work fills a conceptual gap in models of neuronal homeostasis, by demonstrating that neurons and other excitable cells do not necessarily require specific sensing pathways for physiological activity. Generic time-averaged readouts of calcium concentration can support sophisticated sensing and control mechanisms when coupled with the appropriate mix of voltage-dependent conductances.

Canonical transcriptional regulators, such as calcium-calmodulin kineses, may therefore serve as general purpose regulators that can be adapted to sense and control specific firing properties in specific cell types [7]. An immediate prediction of this conceptual model is that targeted deletions of channels should upset homeostatic sensing. This prediction runs counter to more conventional accounts of homeostasis, which predict that channel deletions should elicit responses that compensate, or partially compensate, the role of the missing channel type. To date, experimental evidence for conventional homeostatic responses is mixed, prompting large scale studies to question the existence of homeostatic mechanisms [52]

Our findings also predict that firing rate statistics are coupled in a cell-specific manner. This arises from the simple mathematical observation that rate statistics are necessarily chained in the Volterra series representation of the neuronal membrane response to input. The fact that the variance of the firing rate shapes higher order statistics in a convolutional way, and hence that the firing rate distribution is constrained from the first two moments up to higher-order moments, has not been previously pointed out to the best of our knowledge (although, for a related discussion see [53] chapter 1.7 and references therein). Other work has studied how neuronal firing rate depends specifically on the correlations in the inputs, assuming point process models that do not account for constraints due to nonlinear membrane potential dynamics [54–56].

The implications for the control of mean and variance of firing rate discussed here can be tied to the generic dual-control system proposed by Cannon and Miller in a series of publications [28, 32, 33]. They analytically demonstrated that hitting a *characteristic* (*i*.*e*. joint) mean and variance in firing rate is necessary for the controller to converge to two targets simultaneously. A conclusion common to this previous work and ours is that there is a constraint on the set of reachable targets. However, in our work this constraint is not imposed to ensure stability, it instead arises naturally because mean-variance coupling is constrained by the conductance distribution.

Our main contribution is showing that neurons have cell-specific sensory modes as a corollary of the fact that the intrinsic neuronal dynamics, namely Calcium influx, make this sensing possible. Although our results are limited to the two behaviourally different model neurons we used, we expect them to be broadly valid for various cell types. Future work in this direction could involve more heterogeneous conductance-based models where a larger diversity of sensing modalities might be queried. It would also be interesting to experimentally test our prediction that sensing modes are cell-type specific in a variety of neurons and, specifically, in how cellular dynamics manipulate Calcium concentration.

## Methods

Two classes of conductance-based models were used to simulate intrinsically active and input-driven neurons and to obtain relationships between firing rate statistics and ⟨[*Ca*]⟩.

### Conductance-based model neurons

Neurons receive an applied input current *I*_*app*_(*t*) and have intrinsic ionic currents of the form *I*_*i*_(*v*) = *g*_*i*_*m*_*i*_(*v*)^*p*^*h*_*i*_(*v*)^*q*^(*v* − *E*_*i*_) in addition to a leak current *I*_*leak*_ = *g*_*leak*_(*v* − *E*_*leak*_). The voltage equation is

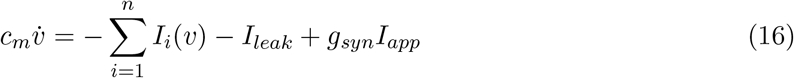

where *c*_*m*_ is membrane capacitance, *g*_*i*_ are maximal conductances, *E*_*i*_ reversal potentials, *m*_*i*_, *h*_*i*_ gating variables and exponents *p, q* ∈ N. Unless otherwise specified, the synaptic conductance *g*_*syn*_ = 1. Neuronal parameters used in simulations are given in table 1. The sum of all Calcium currents, 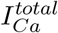, is translated into concentration [*Ca*], in *µM*, through

**Table 1.** Default values for model neurons.

| Variable | Value | Units |
| --- | --- | --- |
| Specific capacitance $c_m$ | 10 | $nF/mm^2$ |
| Calcium buffering time constant $\tau_{Ca}$ | 20 | $ms$ |
| Calcium averaging time constant $\tau_{av}$ | 6000 | $ms$ |

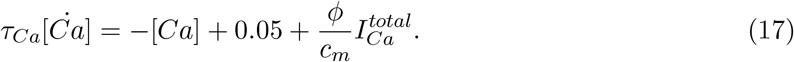

The factor 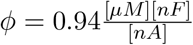 that multiplies 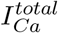 is given for a particular surface area of 0.0628 *mm*^2^ and, thus, for a capacitance *C* = 0.628 *nF*. After filtering with

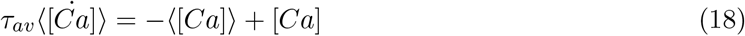

to smooth out the fast fluctuations of [*Ca*], the running average state is ⟨[*Ca*]⟩. The value of this state at the end of the simulated time *T* = 3 × *τ*_*av*_ = 18 *s* gives a scalar measure of average [*Ca*].

In the stomatogastric ganglion (STG) model of crabs [25], there are *n* = 7 ionic currents: a fast sodium *I*_*Na*_; a delayed rectifier *I*_*Kd*_, a fast transient *I*_*Ka*_, and a Ca-activated *I*_*KCa*_ potassium currents; two calcium currents, a fast transient *I*_*CaT*_ and a slow *I*_*CaS*_; and a hyperpolarizationactivated inward cation current *I*_*H*_. A population of 1080 intrinsically active STG neurons is generated through random search in a hypercube. The 4 ionic conductances that are randomized within certain ranges, see table 2, are all scaled by *g*_*leak*_*/*0.01. The values of *g*_*Na*_, *g*_*Ka*_ and *g*_*H*_ are found with a compensation algorithm that uses dynamic input conductances to solve a linear system of equations and produces bursting or tonic-spiking firing patterns [57]. This model has a dynamic reversal potential 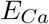 computed from the intracellular Calcium concentration using the Nernst equation with an external Calcium concentration of 3 *mM*; the implementation follows [25].

**Table 2.** STG biophysical parameters.

| Variable | Value | Units |
| --- | --- | --- |
| $g_{CaS}$ | [6, 35] | $\mu S/mm^2$ |
| $g_{CaT}$ | [1, 10] | |
| $g_{KCa}$ | [30, 600] | |
| $g_{Kd}$ | [50, 150] | |
| $g_{leak}$ | [0.02, 0.07] | |
| $g_{Na}$ | * | |
| $g_{Ka}$ | * | |
| $g_H$ | * | |
| $E_{Ca}$ | * | $mV$ |
| $E_K$ | -80 | |
| $E_{leak}$ | -50 | |
| $E_{Na}$ | 50 | |
| $E_H$ | -20 | |

**Table 3.** Biophysical parameters for input-driven neurons.

| Variable | Value | Units |
| --- | --- | --- |
| $g_{Na}$ | [140, 600] | $\mu S/mm^2$ |
| $g_{Ca}$ | [0, 2] | |
| $g_{Ka}$ | [0, 210] | |
| $g_{Kd}$ | 20 | |
| $g_{leak}$ | 0.3 | |
| $E_{Na}$ | 55 | $mV$ |
| $E_{Ca}$ | 120 | |
| $E_K$ | -75 | |
| $E_{leak}$ | -17 | |

Initially, baseline statistics of firing rate are evaluated across all neurons with minimal noise (*σ*_*app*_ = 0.1) and confirm various activity regimes: the mean of the instantaneous firing rate ranges between 0.85 and 150 *Hz*. Baseline variance is zero across the tonic subpopulation on account of their regular firing. For bursters, however, it can reach up to 7 ×10^3^ *Hz*^2^ because the instantaneous rate within bursts can be much higher than the instantaneous rate outside of a burst.

A different model is used to simulate input-driven neurons in sensory brain areas. For this, the Connor-Stevens model is adapted by adding a calcium current following Drion et. al. [58]. There are *n* = 4 ionic currents: a transient sodium current *I*_*Na*_, a delayed rectifier *I*_*Kd*_ and fast transient *I*_*Ka*_ potassium currents, and a non-inactivating calcium current *I*_*Ca*_. To generate the input-driven population of 809 neurons, three conductances are randomly selected in a hypercube while the leak and delayed potassium conductances are fixed. Trials that exhibit bistability in firing are discarded since depolarization blocks shunt input and hence do not represent a truly input-driven neuron. Across the input variance ramping protocol, firing rates are capped at 120 Hz and variance is also capped at 2 × 10^3^ Hz^2^, consistent with the fact that this model produces fast tonic spiking.

### Input and output statistics

The input *I*_*app*_(*t*) fed in to equation 16 as a precomputed time series is pink noise with mean *µ*_*app*_ in units of *nA* and variance 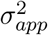. The signal is generated by filtering white noise with a 1*/f* frequency response and has a cut-off frequency of 200 Hz. This represents incoming brain signals that have been filtered through various stages of neuronal processing. Fifteen different instances of *I*_*app*_(*t*) are created with the same statistics to compare results across 15 trials for each input condition: *σ*_*app*_ ∈ [1, 14] and fixed mean, or, *µ*_*app*_ ∈ [1, 10] and fixed variance.

Given an input condition 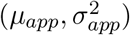, voltage and Calcium trajectories are recorded to extract output statistics and average Calcium concentration. The inter-spike interval

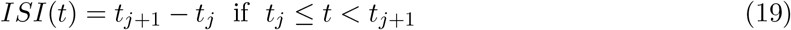

is used to compute the instantaneous firing rate, *r*(*t*) = 1*/ISI*(*t*). Mean firing rate is

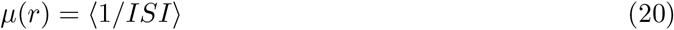

where ⟨·⟩ is the mean of the collection 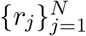. The variance of the firing rate is

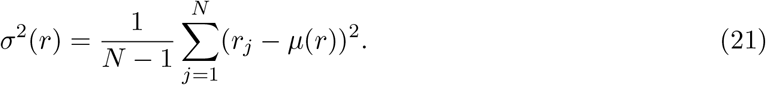

Rolling rate statistics are computed over a window of size *w* = 12*T* (= 216 s) when the controller equations are implemented over a long time course; *τ*_*u*_ = 20 × *τ*_*av*_.

## Acknowledgements

A. R.-H. funded by Huawei Graduate Studies Student Funding. Thanks Dr. Kristine Heiney and Dr. Charles Micou for their helpful comments and valuable feedback on an earlier version of the manuscript.

